# The contribution of maternal oral, vaginal, and gut microbiota to the developing offspring gut

**DOI:** 10.1101/2022.12.19.521059

**Authors:** Amber L. Russell, Erin Donovan, Nicole Seilhamer, Melissa Siegrist, Craig L. Franklin, Aaron C. Ericsson

## Abstract

There is limited understanding of how the microbiota colonizing various maternal tissues contribute to the development of the neonatal gut microbiota (GM). To determine the contribution of various maternal microbiotic sites to the offspring microbiota in the upper and lower gastrointestinal tract (GIT) during early life, litters of mice were sacrificed at 7, 9, 10, 11, 12, 14, and 21 days of age, and fecal and ileal samples were collected. Dams were euthanized alongside their pups, and oral, vaginal, ileal, and fecal samples were collected. This was done in parallel using mice with either a low-richness or high-richness microbiota to assess the consistency of findings across multiple microbial compositions. Sample were analyzed using 16S rRNA amplicon sequencing. The similarities between matched pup and dam samples were used to determine the contribution of each maternal source to the pup fecal and ileal composition at each timepoint. As expected, similarity between pup and maternal feces increased significantly over time. During earlier time-points however, the offspring fecal and ileal microbiotas were closer in composition to the maternal oral microbiota than other maternal sites. Prominent taxa contributed by the maternal oral microbiota to the neonatal gut microbiota were supplier-dependent and included *Lactobacillus* spp., *Streptococcus* spp., and a member of the *Pasteurellaceae* family. These findings align with the microbial taxa reported in infant microbiotas, highlighting the translatability of mouse models in this regard, as well as the dynamic nature of the gut microbiota during early life.

## INTRODUCTION

The maturation process of the gut microbiota (GM) is an essential process for life-long health that is defined by the acquisition and colonization of microorganisms in the gut and the subsequent immune system induction that occurs during early life. While emerging evidence suggests that initial colonization of the gut may happen as early as *in utero*^1,2^, this is controversial, and the conventional understanding is that the first bacterial seeding of the gut begins at birth. Regardless of the timing, the initial bacterial colonization of the neonatal gut undoubtedly derives from a maternal source, emphasizing the importance of understanding how vertical transfer is initiated and influenced during continued exposure to maternal microbial sources over the course of GM maturation^3,4^. During the process of maturation, the neonatal GM is especially susceptible to environmental factors capable of inducing persistent effects on the developing GM^5^. Previously identified factors within both human and mouse model populations associated with significant effects on the composition of the neonatal microbiota include mode of delivery^6,7^, breastfeeding^8,9^, and antibiotic exposure^10,11^. These factors have the potential to confound research in human neonates, making characterization of the development and contribution of maternal sources to the neonatal gut microbiota difficult^5^.

There is strong agreement across host species regarding the basic characteristics of normal GM maturation that correspond with physiological changes occurring in the gastrointestinal tract^12,13^. During early life, the neonatal gut microbiota of vaginally born, breastfed infants and nursing mouse pups alike is dominated by members of the *Enterobacteriaceae* and *Lactobacillaceae* families^4,14^, which are transferred from mother during parturition and subsequently colonize the neonatal GM due to the high oxygen availability in the gastrointestinal tract relative to adults^12,15^. Next, as lumenal oxygen tension decreases, and breastfeeding continues, *Bifidobacteriaceae* and *Clostridiaceae* proliferate until weaning, presumably due to their role in metabolism of milk oligosaccharides and facultative anaerobic capabilities^16,17^. With the introduction of solid foods^13^, a more diverse and adult-like gut microbiota develops, with *Bacteroidaceae, Lachnospiraceae*, and *Ruminococcaceae* being dominant families of the mature gut microbiota^5,13,18^, and the maturation process can be considered complete once the GM has reached a richness and composition consistent with that of the maternal feces.

There is limited knowledge regarding the contribution of different maternal source microbiotas in the neonatal mouse gut microbiota. Literature describing the vertical transfer and maturation of the human gut microbiota has relied on noninvasive sampling, and largely ignored the upper gastrointestinal tract (GIT)^3,4,19^. The few studies of the developing murine gut microbiota that included samples of the upper GIT found compositional and functional differences compared to the lower GIT, but the maternal source of ileal microbes was not investigated^20,21^.

To address these knowledge gaps, we characterized the neonatal fecal and ileal microbiota at multiple pre-weaning time-points, alongside the maternal fecal, ileal, vaginal, and oral microbiotas using targeted amplicon sequencing. Core microbiota analyses and similarities between matched neonatal and maternal samples were used to assess the relative contribution of the different maternal sites to the developing neonatal microbiota in both compartments of the GIT. Additionally, considering the significant differences in richness and beta-diversity among SPF microbiotas from different producers, the entire experimental design was replicated in isogenic mice with microbiotas originating from two different U.S. suppliers^22^, The Jackson Laboratory and Envigo.

## RESULTS

### Phylum-level composition changes rapidly during gut microbiota development

Neonate feces demonstrated incrementally lower relative abundance (RA) of *Bacillota* and higher RA of *Bacteroidota* at each subsequent timepoint (Figure 1A, B), with the latter proliferating substantially on day 11 or 12 of life. Prior to day 12, high richness GM4 neonate feces also contained high RA of *Pseudomonadota*, while this phylum was present at a much lower RA in low richness GM1. The same difference was observed in the neonate ileum (Supplemental Figure 1A, B). Maternal samples from GM1 (Figure 1C) also contained lower proportions of *Pseudomonadota* than maternal samples from GM4 (Figure 1E), while also harboring a greater number of bacterial phyla within vaginal and fecal samples. The vaginal and fecal microbiota of GM4 maternal samples contained populations of *Bacillota, Bacteroidota, Deferribacterota, Verrucomicrobiota, Cyanobacteria*, and *Actinomycetota* at proportions of 1% or greater, while GM1 samples contained only *Bacillota, Bacteroidota*, and *Pseudomonadota* at proportions of 1% or greater.

**Figure 1.**
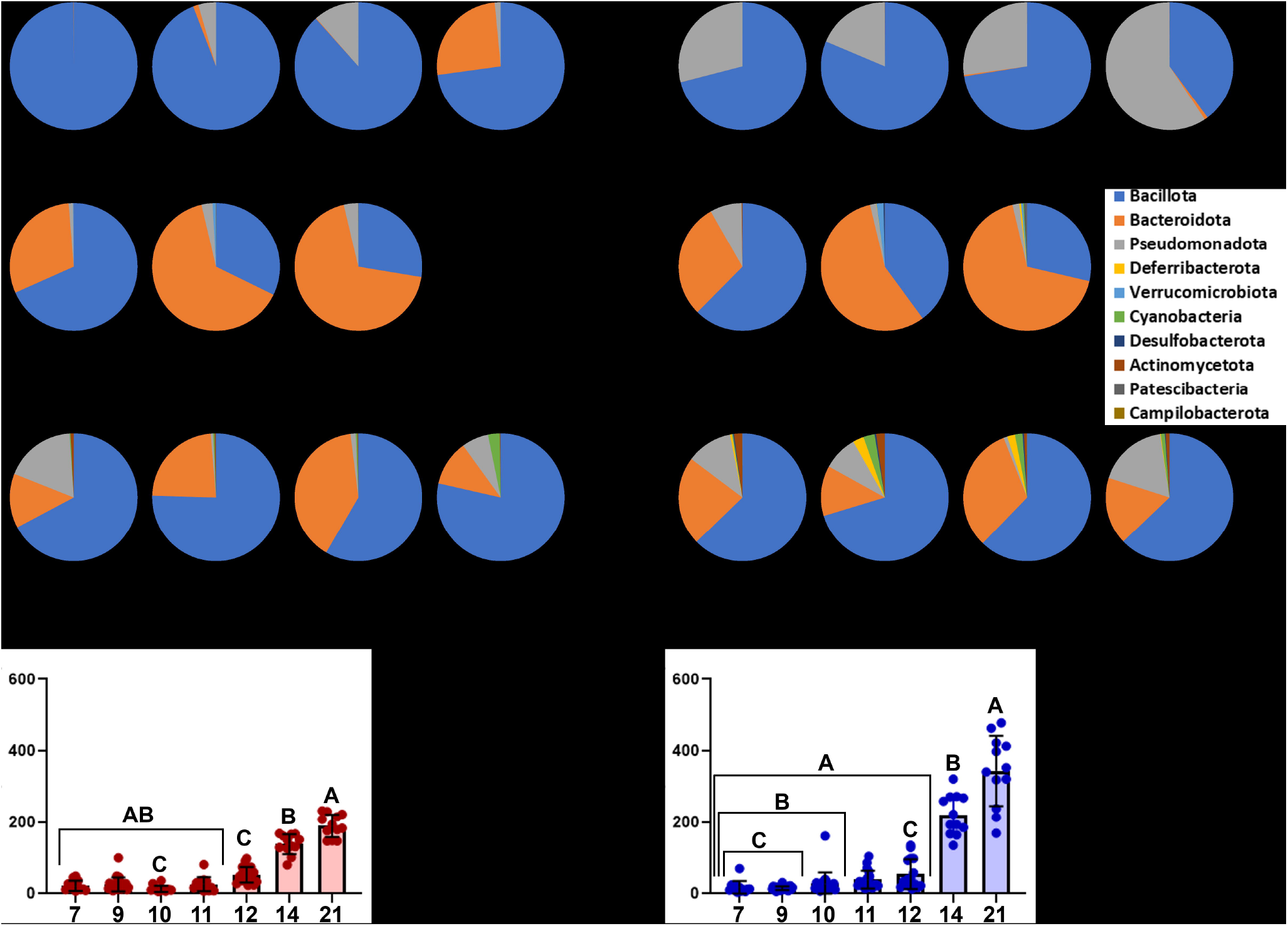
Fecal Bacteroidota relative abundance and alpha diversity are greater in older pups. (a-d) Circle graphs depicting average relative abundance of samples. Figure key located on right side of panel. (a-b) Pup fecal samples grouped by timepoint from (a) GM1 and (b) GM4 mice. (c-d) Maternal tissue samples groups by sample tissue type for (c) GM1 and (d) GM4 mice. (e-f) Bar charts depicting alpha diversity of samples using Chao-1 index with bars representing group averages and error bars representing standard error of mean (SEM). Each symbol represents an individual sample. (e) Average GM1 pup fecal alpha diversity grouped by timepoint. (f) Average GM4 pup fecal alpha diversity grouped by timepoint. Statistics were calculated using ANOVA on ranks within each GM, like letters within a graph denote a significant difference in pair wise comparisons using Dun’s post-hoc.

An ANOVA on ranks test was used to identify differences in richness within each GM, using the Chao-1 index. As expected, the richness of neonate fecal samples was significantly higher at later timepoints, however within ileal samples, age did not have a significant main effect on richness in either GM. For GM1 mice, fecal richness at day 14 and day 21 was significantly higher than samples from day 7 through day 11 (Figure 1E), while ileal richness had no clear pattern based on timepoint (Supplemental figure 1). A similar effect was observed in GM4 neonate richness, where fecal richness at day 21 was significantly higher than day 7 through 12 (Figure 1F), and no difference in richness was detected in the ileum (Supplemental figure 1).

### Evidence of GM-specific vertical transfer

To better resolve the taxonomic differences between GM1 and GM4 pups within the *Bacillota* and *Pseudomonadota* phyla, we determined the RA of ASVs annotated to the order *Lactobacillales* and the phylum *Pseudomonadota*. The RA of *Lactobacillales* was lower at day 21 compared to day 7 through day 12 in both GM1 and GM4 neonate fecal samples (Figure 2A). Regarding the different maternal sites (Figure 2B), the RA of *Lactobacillales* was variable in both GMs, with no statistical difference between sampling sites for either GM1 (H= 17.91, *p* = 0.001) or GM4 (H = 7.34, *p* = 0.12). The oral microbiota contained GM-specific species of *Streptococcus*, including *S. danieliae* in GM1, and *S. merionis* in GM4, and this difference was reflected in the neonatal feces. Within the *Pseudomonadota*, an ASV resolved to the family *Pasteurellaceae* was dominant in both GMs, particularly in GM4, reaching a mean RA greater than 50% at day 11 before declining significantly at all later time-points (Figure 2C). A similar trend was observed in *Escherichia-Shigella*, wherein the RA was higher within GM4 at day 12 compared to earlier timepoints (day 7: *p* = 0.04, day 9: *p* = 0.007). On day 14 and 21, *Pasteurellaceae* and *Escherichia-Shigella* were largely replaced by *Parasutterella excrementihominis* in GM1, and an unspeciated *Parasutterella* in GM4 (Figure 2C). A high RA of the dominant *Pasteurellaceae* ASV was detected in maternal oral samples, but this difference did not reach significance compared to other maternal sites.

**Figure 2.**
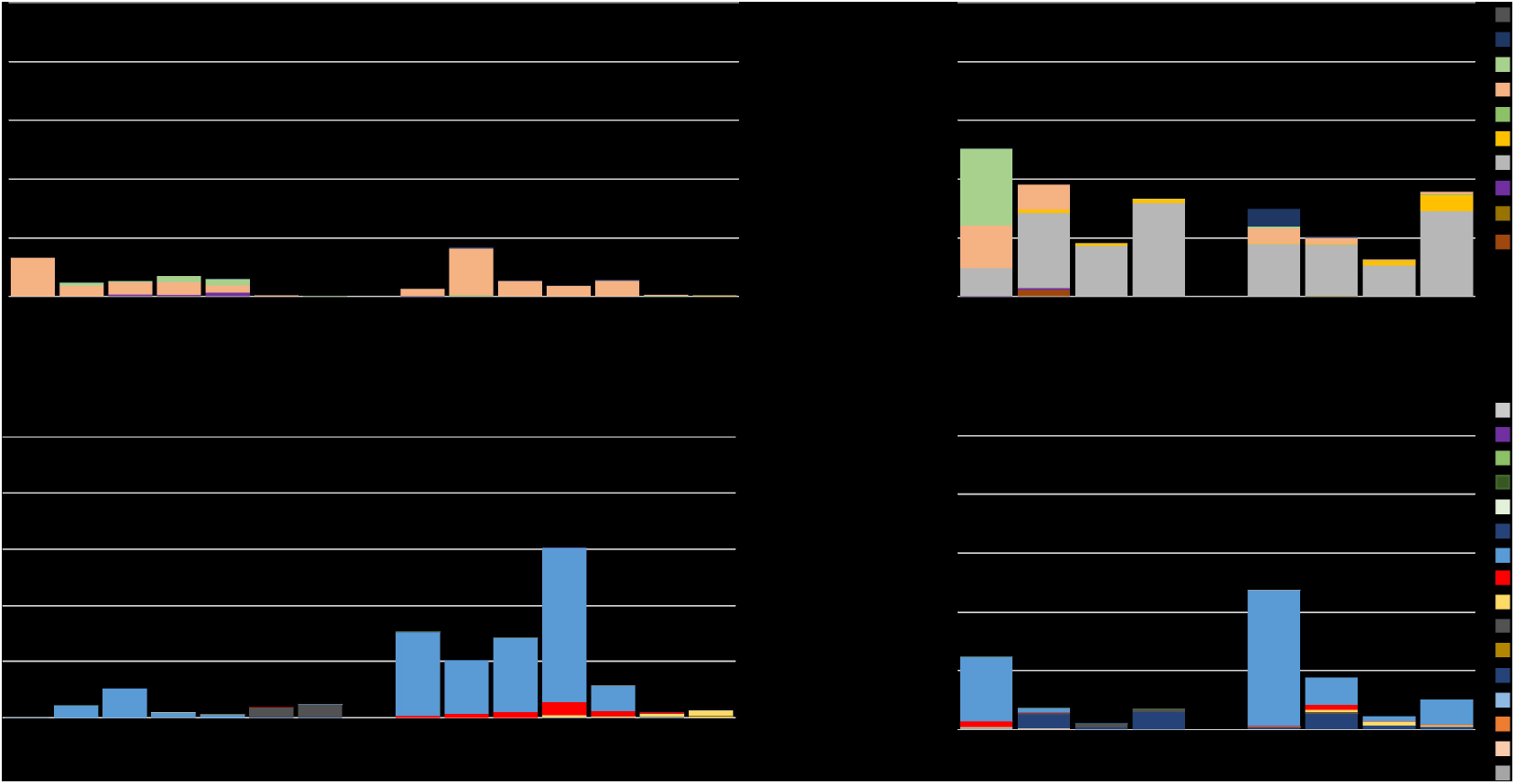
GM4 Pup feces harbors a higher proportion of *Pseudomonadota* species than GM1. (a-b) Bar charts depicting the average proportion of taxa annotated to *Lactobacillales* in study samples. Figure key is located on the right side of panel. (a) pup fecal samples grouped by GM and age of pups. (b) Maternal samples grouped by GM and tissue type. (c-d) Bar charts depicting the average proportion of ASVs annotated to *Pseudomonadota* in study samples. Bar charts depicting the average proportion of taxa annotated to *Pseudomonadota* in study samples. Figure key is located on the right side of panel. (c) pup fecal samples grouped by GM and age of pups. (d) Maternal samples grouped by GM and tissue type.

The neonatal ileum was distinct from feces but also reflected several trends seen in the neonate feces, including GM1-specific growth of *S. danieliae* (Supplement Figure 2). Similarly, *Pasteurellaceae* represented the dominant *Pseudomonadota* within the neonatal ileum, showing a similar peak RA approaching 50% in the GM4 neonate ileum at day 11 (Supplemental Figure 2). Collectively, these data suggest that dominant taxonomies are shared between the maternal oral cavity and neonate mouse gut.

### Similarity to maternal tissue depends on neonatal GI sample location

Principal coordinate analysis (PCoA) was used to visualize the beta diversity of neonatal and maternal samples within each GM. Comparison of all maternal sites to neonate feces at the earlier timepoints indicated a greater similarity between oral samples and neonate feces than other maternal sites, in both GM1 (Figure 3A) and GM4 (Figure 3B).

**Figure 3.**
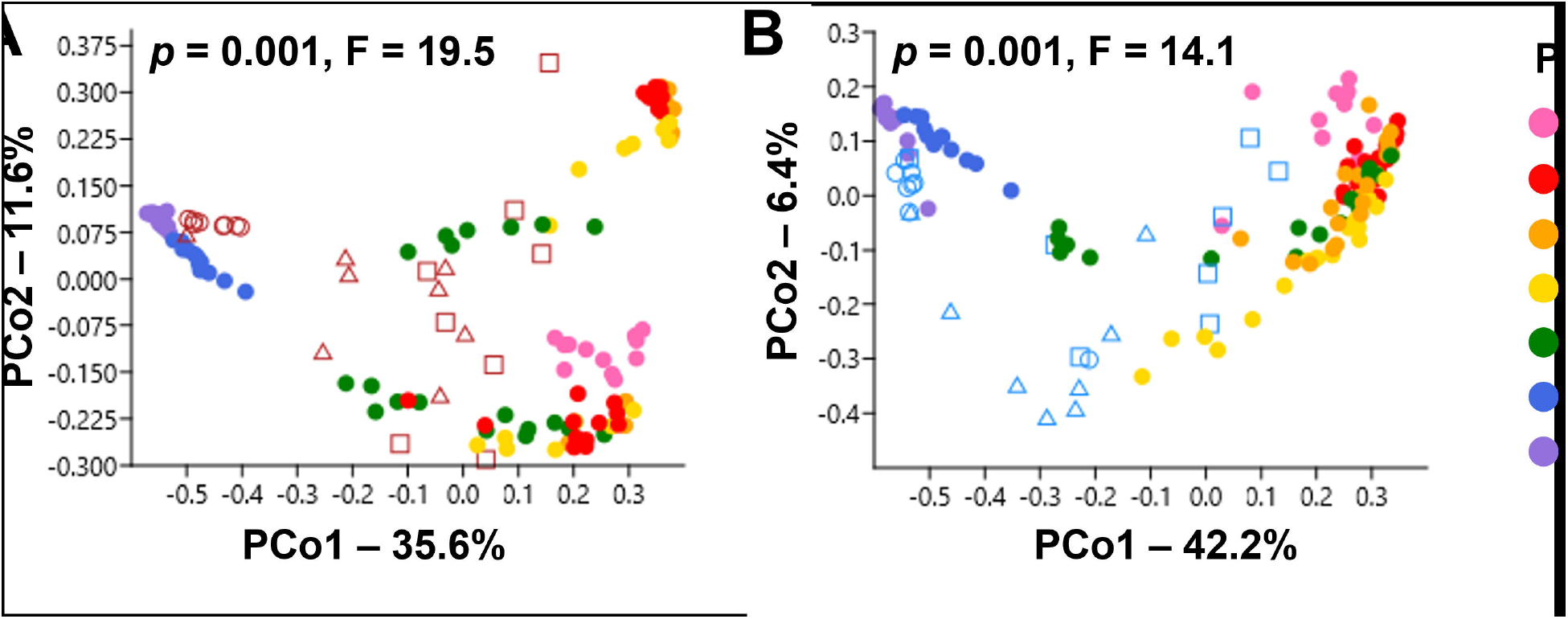
Neonatal fecal similarity to maternal tissues shifts dramatically with time in both GMs. Principal coordinate analysis (PCoA) graphs depicting beta diversity using Bray-Curtis similarity index. Figure key to the right side of panel. (A-B) Pup fecal samples with maternal oral, vaginal, and fecal samples from groups (a) GM1 (b) GM4. Statistics were calculated using a one-factor PERMANOVA for each graph.

Within ileal samples, there was less clustering according to timepoint, and less separation from maternal samples. Within GM1 and GM4 (Supplemental Figure 3), maternal oral samples clustered more closely to the neonate ileal samples than did the other maternal sites.

Collectively, these data suggest that the offspring gut microbiota does not mature until the third week of life, and that the immature gut microbiota is more similar in composition to the maternal oral microbiota than the vaginal microbiota.

### The maternal vaginal contribution to neonatal SPF microbiota is negligible

To control for dam-to-dam variation, we also calculated the Bray-Curtis similarity between each pup fecal sample and matched maternal tissue samples. A three-way ANOVA was used to identify main effects and interactions between GM, age of pups, and maternal sampling site.

There was a significant main effect of all factors, and significant interactions between all three factors analyzed (F = 2.38, *p* = 0.005). In order to analyze pairwise comparison between samples within each GM, data were stratified by GM and a two-way ANOVA was utilized. Within GM1, neonate feces had the highest similarity to maternal fecal samples at all timepoints compared to oral and vaginal samples (Figure 4A). The similarity between pup feces and the maternal oral and vaginal microbiotas was much lower and favored similarity to the oral microbiota during early life. Within GM4, maternal oral samples had significantly higher similarity to pup feces than maternal vaginal or fecal samples from day 7 to day 12 (Figure 4B). By day 14, similarity of offspring feces to maternal feces was significantly higher than similarity to oral (*p* < 0.001, t = 12.48) or vaginal samples (*p* < 0.001, t = 14.64), which remained true on day 21.

**Figure 4.**
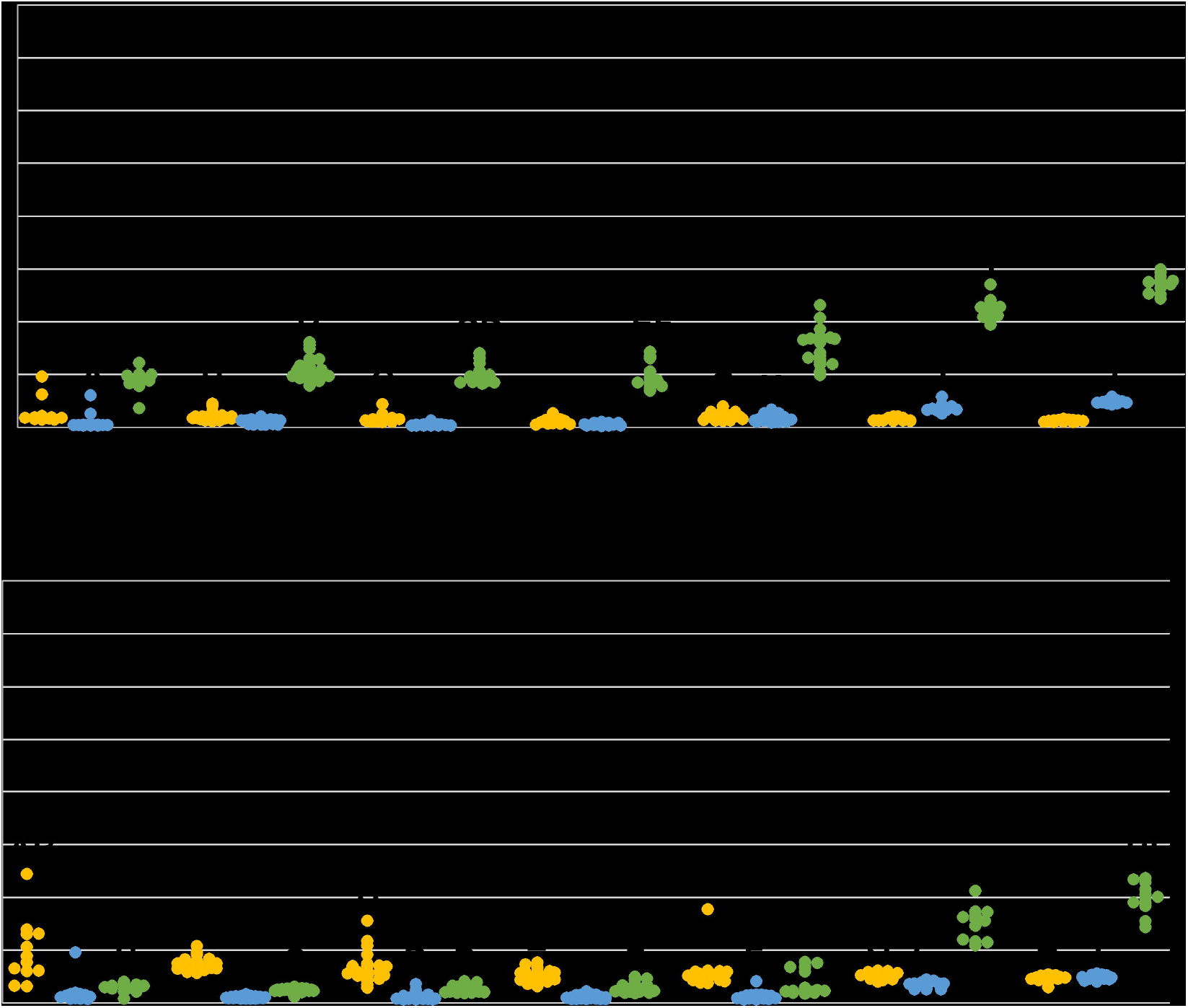
Fecal similarity increases over time, but similarity to maternal oral microbiome is GM dependent. (a-b) Dot plots and bar graphs depicting Bray-Curtis similarity of pup feces to maternal tissue. (a) Average Bray-Curtis similarity of maternal samples to pup feces in GM1. Each dot represent the average bray-Curtis similarity of one pup fecal sample to all maternal samples of that tissue type in GM1. (b) Average Bray-Curtis similarity of maternal samples to pup feces in GM4. Each dot represent the average bray-Curtis similarity of one pup fecal sample to all maternal samples of that tissue type in GM4. Statistics were calculated within each GM using a Two-way ANOVA on tissue type and age of pups. Matching letters denote significant pairwise differences detected during a Holm-Sidak post-HOC analysis. O-Oral, V-Vaginal, F-Fecal.

Similarities between ileal samples and matched maternal samples were also compared, during the period between days 9 and 12. A three-way ANOVA found an overall main effect of age of pups (F = 12.30, *p* < 0.001) and maternal tissue (F = 13.48, *p* < 0.001), as well as interactions between all three factors (F = 6.20, *p* < 0.001). Two-way ANOVA stratified by GM found a significant difference between maternal tissue type within GM1 on day 10 and 11 (Supplemental Figure 4). Within GM4 ileal samples, there was a significant difference between maternal tissue type at each timepoint. While similarity to maternal sites in each GM varied from day to day during this period, the highest similarity was most often to maternal oral or ileal samples.

Next, we utilized Spearman correlation analyses to determine the correlation between maternal tissue similarity to both the neonatal upper and lower GIT. Neonates that had both fecal and ileal samples collected were used for this analysis (n = 146). There was a significant positive association between maternal oral and neonatal GIT similarity (Supplemental Figure 4). Maternal vaginal similarity to neonatal fecal and ileal samples did not correlate significantly.

Neither maternal vaginal nor maternal fecal samples correlated with neonatal GIT samples in terms of similarity. These data suggest that those mice in which the maternal oral microbiota is most represented in the neonatal ileum, are expected to have the highest similarity between maternal oral and neonatal fecal microbiota as well.

### Compositional similarity of the neonatal GIT to the maternal feces increases with age

To determine the number of ASVs consistently shared between the neonatal fecal microbiota and maternal tissue sites over the course of gut microbiota development, the core microbiota^23^ of each maternal tissue within each GM was compared to the core microbiota of neonatal feces at each timepoint within the same GM. An ASV was considered part of the core microbiota if it was detected in 30% or more of samples within a group and had an average group relative abundance of 0.01% or higher. Over the course of fecal microbiota development, the core microbiota of GM1 neonates first undergoes an expansion of ASVs exclusively found in the oral maternal microbiota (Figure 5A). This event consists of four ASVs detected from day 9 to day 12. The second expansion involves ASVs shared exclusively with the maternal fecal core microbiota starting at day 11. With each consecutive timepoint, the contribution of maternal core fecal ASVs increased up to the final timepoint, with 79 ASVs detected (Figure 5A). For the neonatal fecal core microbiota in GM4, ASVs shared exclusively with maternal fecal samples were not detected until day 12 with nine ASVs, which increased to 66 ASVs by day 21 (Figure 5B). The majority of core ASVs detected in GM4 were not exclusive to any maternal tissue core.

**Figure 5.**
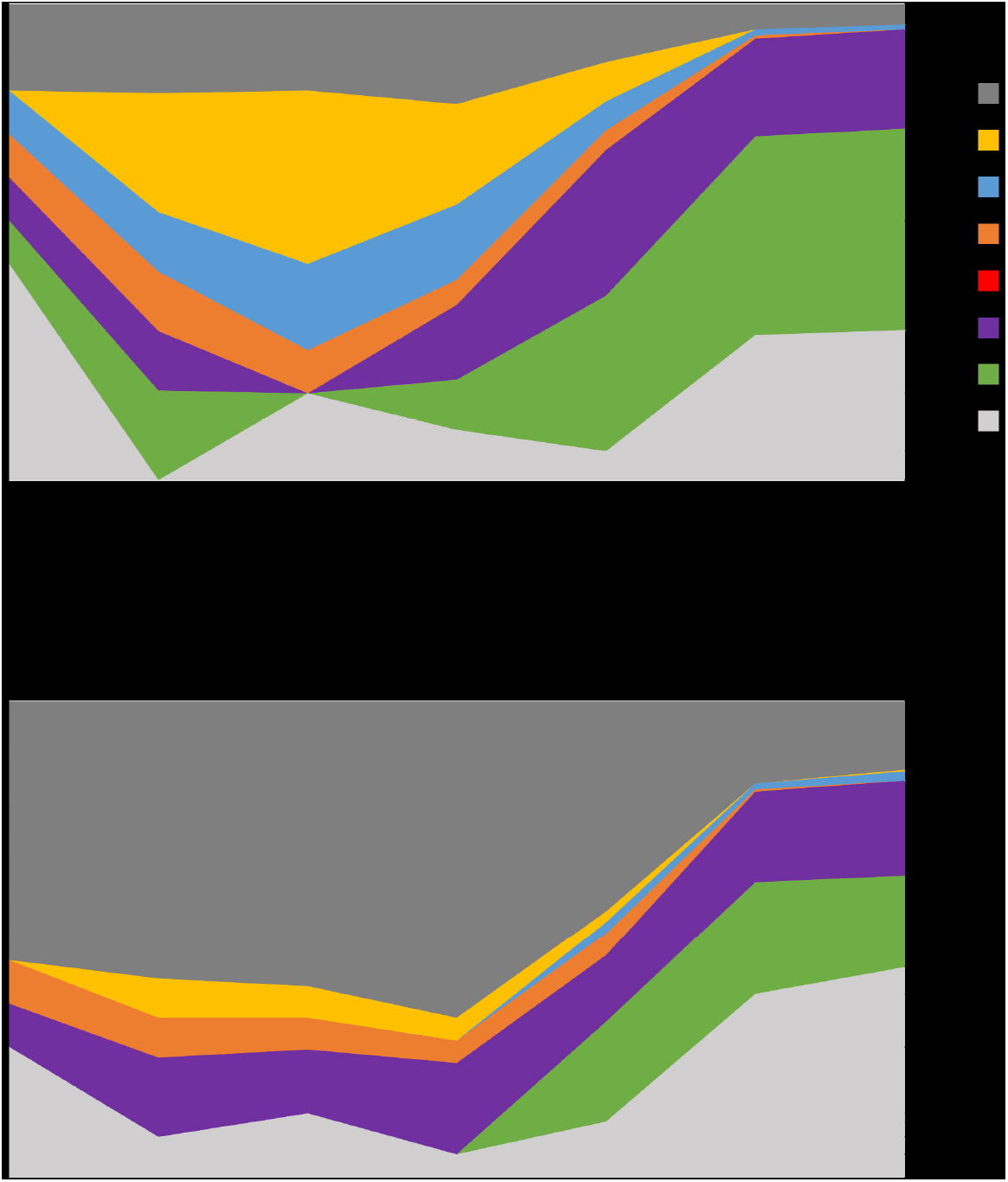
Unique maternal vaginal ASVs were not incorporated into pup fecal microbiota. Area graphs depicting the proportion of amplicon sequence variants (ASV)s found in the pup fecal core microbiota at each timepoint, that are also found in maternal tissue core microbiota. Figure key is located to the right of the panel. (a) GM1 pup fecal and (b) GM4 pup fecal core microbiota.

The same approach for identifying core microbiota was used for ileal samples. For the GM1 ileal core microbiota, there was only one ASV shared exclusively with the maternal ileal core microbiota at any given timepoint, while the neonatal ileal core microbiota shared up to seven ASVS exclusively with the maternal oral core microbiota at days 9, 10, and 11 (Supplemental Figure 5). The GM1 neonatal ileal core microbiota was the only neonatal core microbiota to share ASVs exclusively with the maternal vaginal core microbiota. The core microbiota of GM4 neonatal ileal samples mostly contained ASVs that were shared between all maternal tissue core microbiotas, and there was only one ASV shared exclusively with one maternal source (oral) at any given time.

## DISCUSSION

Our results indicate that the gut microbiota of neonatal mice rapidly changes during development to approach the maternal gut microbiota in composition and diversity by three weeks of age (Figure 1). Within feces, proportions of *Bacillota* and *Pseudomonadota* decrease, while proportions of *Bacteroidota* increase throughout maturation. These changes in phylum level RA are consistent with previous findings indicative of the diversification of the gut microbiota population seen in both humans and mice at weaning^24^. The expansion of *Bacteroidota* is correlated with the onset of consumption of plant derived carbohydrates found in solid foods in both humans and mice^13^, likely explaining the increase seen in this study. The decline in the proportion of *Pseudomonadota* and *Bacillota* as pups age may be due to the shift from an aerobic to an anaerobic environment within the intestines due to the decline in oxygen availability as pups age, causing a compositional shift favoring bacteria that are able to utilize anerobic respiration^25^.The maturation of the gut microbiota is further exemplified in these pups by the steady increase in fecal alpha diversity over time, as has been reported previously during early development in both species^4,5,26,27^.

The RA of the phylum *Pseudomonadota* was strikingly different between GM1 and GM4 in both neonatal fecal and ileal samples (Figure 2, Supplemental Figure 2). Specifically, GM4 mice harbored higher proportions of *Pseudomonadota* compared to GM1 mice, as two ASVs annotated to *Pasteurellaceae* and one *Escherichia-Shigella* ASV were enriched in GM4 compared to GM1 neonatal feces. Interestingly, maternal *Escherichia-Shigella* was found in highest proportions in oral samples in GM1 but in vaginal samples in GM4. We also found that the RA of *Pseudomonadota* changes in the neonatal gut microbiota over time, but *Lactobacillales* colonization appears less dynamic during the maturation of the gut microbiota. This may be due to intermittent presence of *Pseudomonadota* in the maternal milk microbiota^9,28^ and the ability of *Pseudomonadota* to metabolize milk oligosaccharides^29^, while the decline in *Pseudomonadota* in neonatal mice has been associated with increased IgA production as neonates age^30^. In contrast, *Lactobacillales* are lactic acid-producing bacteria that utilize carbohydrates and colonize the gut throughout infancy and adulthood and can survive in both anerobic and aerobic conditions^31^. The higher proportions of *Streptococcus danieliae* in GM1 and *Streptococcus merionis* in GM4 in the neonatal gut microbiota were vertically transferred from the maternal oral microbiota. These results suggest that different background GMs are predisposed to harboring not only different proportions of bacteria, but also experience species-specific gut microbiota vertical transfer of these populations.

As pups age, both fecal and ileal GM beta diversity became increasingly similar to maternal fecal and ileal samples respectively, as well as inter-individual similarity within pups (Figure 3 and Supplemental Figure 3). This increase in similarity is expected, and has repeatedly been reported in the literature of both human and mouse model as a hallmark of GM^14,32^. The contribution of maternal microbial sites to the neonatal gut microbiota varies between different SPF GMs. In terms of beta diversity, the similarity of offspring feces to oral maternal samples remained stable overtime, however in GM4, this similarity was significantly higher than fecal maternal similarity until day 14. This provides evidence that maternal tissue contribution varies between gut microbiota compositions, and we may thus infer that vertical transfer from maternal microbial sites is mediated in part by the composition of the various source microbiotas. The similarity of offspring ileum to maternal tissue samples was inconsistent over time, but findings suggest the same trends seen in the feces.

Analysis of beta diversity similarity (Figure 4, Supplemental Figure 4) and the core microbiota of pup samples in relation to the maternal oral, vaginal, fecal, and ileal composition (Figure 5, Supplemental Figure 5) revealed high similarity between pup feces and maternal oral samples in early life, with increasing similarity to maternal gastrointestinal samples as pups age. These results are consistent with recent studies, in which maternal vaginal samples do not colonize the neonatal gut microbiota as well as other maternal sources^3^. As such, this study provides further evidence of conserved events between humans and mice during the maternal transfer of microbes and maturation of the offspring gut microbiota.

Mice fecal samples from each GM were originally collected at day 7, 14, and 21 to examine changes in the fecal microbiota overtime. We found a large compositional shift between day 7 and day 14 in which the neonatal mouse fecal microbiota underwent a dramatic change in composition. To examine this compositional shift in greater detail, we focused on days 9, 10, 11 and 12 to examine the bacterial composition of pup ileum in relation to different maternal sites during this period of transition. Although we do not have ileal data for all timepoints, we found the data pertinent to include in this analysis for a more complete understanding of neonatal gut microbiota development.

The results of this study are overall consistent with recent literature regarding gut microbiota development and vertical transfer of various maternal microbiota sources. Previous maternal transfer studies have focused on human rather than mice, as such our study is the first of its kind to analyze the effect of multiple microbiota sources on gut microbiota development in mice. By utilizing the differing compositions of GM1 and GM4, this study was able to model some of the GM variation seen between individuals in human studies, thus allowing us to find significant differences in the contribution of various maternal bacterial on the pup GM composition throughout development. These results support the use of the mouse as an appropriate model for gut microbiota development and highlight the importance of utilizing a study design with more than one gut microbiota composition.

In conclusion, SPF mouse microbiotas undergo a dynamic and somewhat characteristic maturation process, culminating by roughly two to three weeks of age. Prior to that, the neonatal gut microbiota is more similar in composition to the maternal oral microbiota, as opposed to the vaginal of fecal microbiotas. Additionally, the maternal source microbiota that is transferred during gut microbiota development is dependent on the specific SPF microbiota. Further studies are needed to expand our knowledge regarding the effect of these developmental exposures on host development, and if additional maternal bacterial sources, such as the skin, contribute significantly to GM development.

## METHODS

### Animals

All experiments described in this manuscript were approved by the Institutional Animal Care and Use Committee (IACUC) of the University of Missouri (protocol 9587). Outbred CD-1 mice maintained at the University of Missouri harboring either a low richness background GM originating from The Jackson Laboratory, designated GM1, or a high richness background GM originating from Envigo, designated GM4. A detailed description of these colonies has been previously published^22^. 16 female mice (GM1: n = 8, GM4: n = 8) were bred and their subsequent pups were allowed to age until either day of age 9 (GM1: n = 23, GM4: n = 22), 10 (GM1: n = 13, GM4: n = 21), 11 (GM1: n = 13, GM4: n = 19), or 12 (GM1: n = 19, GM4: n = 16).

Two dams and their subsequent pups were euthanized at each timepoint chosen at random, and samples were collected immediately after euthanasia. An additional 12 pup fecal samples were also collected at 7, 14, and 21, days of age from colony pups from both GM backgrounds for comparison to the dam-pup matched samples. Mice were housed at Discovery Ridge in Columbia, MO under barrier condition in microisolator cages on Thoren ventilated racks under a 14:10 light/dark cycle. Each cage contained pelleted paper bedding with nestlets and received *ad libitum* access to irradiated Breeder diet 5053 rodent chow and acidified, autoclaved water.

### Sample collection

All maternal samples were collected *post mortem* at days 9, 10, 11, and 12. Oral and vaginal swabs were used to collected oral and vaginal microbiota samples respectively, by inserting a cotton swab into the designated orifice and rotating the swab. Swab samples were then collected into 2 mL round-bottom tubes. Maternal ileal samples were collected by excising roughly 4 cm of ileum proximal to the ileocecal junction and rinsing the luminal contents of the sample into a 2 mL round-bottom tube using sterile PBS. For maternal fecal samples, the two most distal fecal pellets in the rectum or distal colon were collected and placed in 2 mL round-bottom tubes with a 0.5 cm-diameter stainless steel ball bearing for homogenization of sample. Pup feces were collected as described for dams at days 7, 9, 10, 11, 12, 14, and 21 of age. Pup ileal samples were collected at days 9, 10, 11, and 12 of age by excising 2 cm of the ileum proximal to the ileocecal junction and collecting into a 2 mL round-bottom tube with a 0.5 cm-diameter stainless steel ball bearing due to the small size of the ileal lumen. All samples were placed on ice immediately following collection and samples were stored in a -80° C freezer until DNA was extracted.

### DNA extraction

All sample tissue DNA was extracted using PowerFecal Pro kits (Qiagen) according to manufacturer’s protocol, with the exception that samples were homogenized using a TissueLyser II (Qiagen) for 10 minutes at 30/sec, in lieu of a vortex adapter as described by PowerFecal Pro kit instructions. DNA yields were quantified by fluorometry via the quant-iT BR dsDNA reagent kits (Invitrogen) and normalized to a consistent concentration and volume prior to submission for downstream processing.

### 16s rRNA library preparation and sequencing

Tissue sample DNA was processed at the University of Missouri Genomics Technology Core. Bacterial 16S rRNA amplicons were constructed via amplification of the V4 region of the 16S rRNA gene with the universal primer set (U515F/806R) and flanked by Illumina standard adapter sequences as in a method previously described elsewhere^33,34^. Dual-indexed forward and reverse primers were used in all sample reactions. PCR was initiated in 50 uL reactions containing 100 ng metagenomic DNA, primers (0.2 uM each), dNTPs (200 uM each), and Phusion high-fidelity DNA polymerase (1U, Thermo Fisher). Amplification parameters used were as followed: 98°C(3 min) + [98°C(15 sec) + 50°C(30 sec) + 72°C(30 sec)] × 25 cycles + 72°C(7 min). Amplicon pools of 5 µL/reaction were combined, thoroughly homogenized, and then purified with addition of Axygen Axyprep MagPCR clean-up to an amplicon volume of 50 µL. Amplicons were then incubated for 15 minutes at room temperature and received multiple washes of 80% ethanol. Post-wash, the dried pellet was resuspended in 32.5 µL EB buffer (Qiagen), incubated at room temperature for 2 minutes, and then placed on a magnetic stand for five minutes. The final amplicon pool was then evaluated using quant-IT HS dsDNA reagent kits and diluted according to the Illumina standard protocol for sequencing of 2×250 bp pairedend reads. Amplicon pools were then sequenced on the MiSeq instrument.

### Informatics analysis

DNA sequences were assembled and annotated at the MU Bioinformatics and Analytics Core. The primer set was designed to match the 5’ end of both the forward and reverse amplicon reads. Cutadapt4 (version 2021.8.0; https://github.com/marcelm/cutadapt) was then used to remove the primer from the 5’ end of the forward read, and then if found, remove the reverse complement of the primer to the reverse read from the forward read. Therefore, a forward read could be trimmed at both ends if the insert were shorter than the length of the amplicon. The same method was then utilized for reverse reads, but with the primers in the opposite roles.

Read pairs were then rejected if one read or the other did not match a 5’ primer, and the allowed error-rate was 0.1. Two passes over each read count were made to ensure removal of the second primer, and a minimal overlap of three bp with the 3’ end of the primer was required for removal.

The QIIME2 DADA2 plugin (version 1.18.0) was utilized to denoise, de-replicate, and count amplicon sequence variants (ASVs). The following parameters were incorporated: 1) forward and reverse reads were truncated to 150 bases, 2) forward and reverse reads with an expected error higher than 2.0 were ignored, and 3) Chimeras were detected using the “consensus” method and then removed. Python version 3.8.10 and Biom version 2.1.10 were used in QIIME2. Taxonomies were assigned to each of the final sequences using the Silva.v138 database, utilizing the classify-sklearn procedure.

### Statistics

To calculate significant changes in alpha diversity and RA of pup fecal and pup ileal samples, we utilized ANOVA on ranks within each GM, due to a lack of normality. Alpha diversities were quantified by the Chao-1 index using PAST software. ANOVA on ranks was performed using SigmaPlot 14.0 with a Dunn’s *post hoc* analysis for pairwise comparisons.

One-way permutational multivariate analysis of variance (PERMANOVA) was used to test for significant differences in beta diversity of samples and provide pair-wise comparisons of betadiversity. PERMANOVA testing was performed using PAST software using Bray-Curtis similarities. The Bray-Curtis similarity of pup fecal and pup ileal samples to maternal tissues were tested for significant differences within each GM using two-way ANOVAs for the factors tissue type and age using SigmaPlot 14.0. A Holm-Sidak *post hoc* analysis was used to determine significant differences by pair-wise comparison of each group within each two-way ANOVA analysis via SigmaPlot 14.0 (Systat Software, Inc, San Jose, CA). Spearman correlations between pup fecal and pup ileal Bray-Curtis similarities values were calculated using SigmaPlot 14.0.

## Data Availability

Authors are currently working on submitting the necessary files for SRA upload.

