## Supplementary figures and images for "The contribution of maternal oral, vaginal, and gut microbiota to the developing offspring gut"

### Supplemental Figure 1

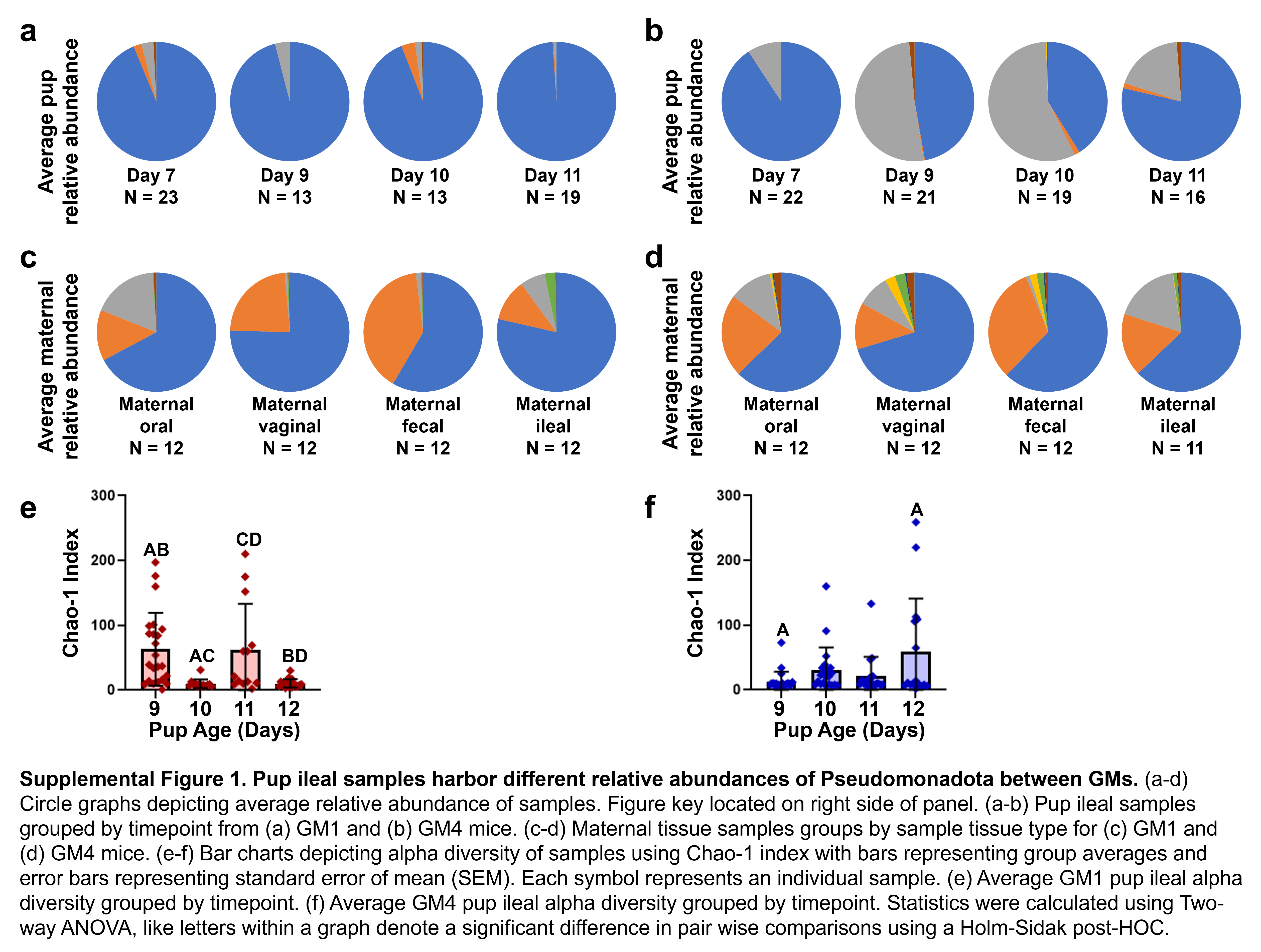

### Supplemental Figure 2

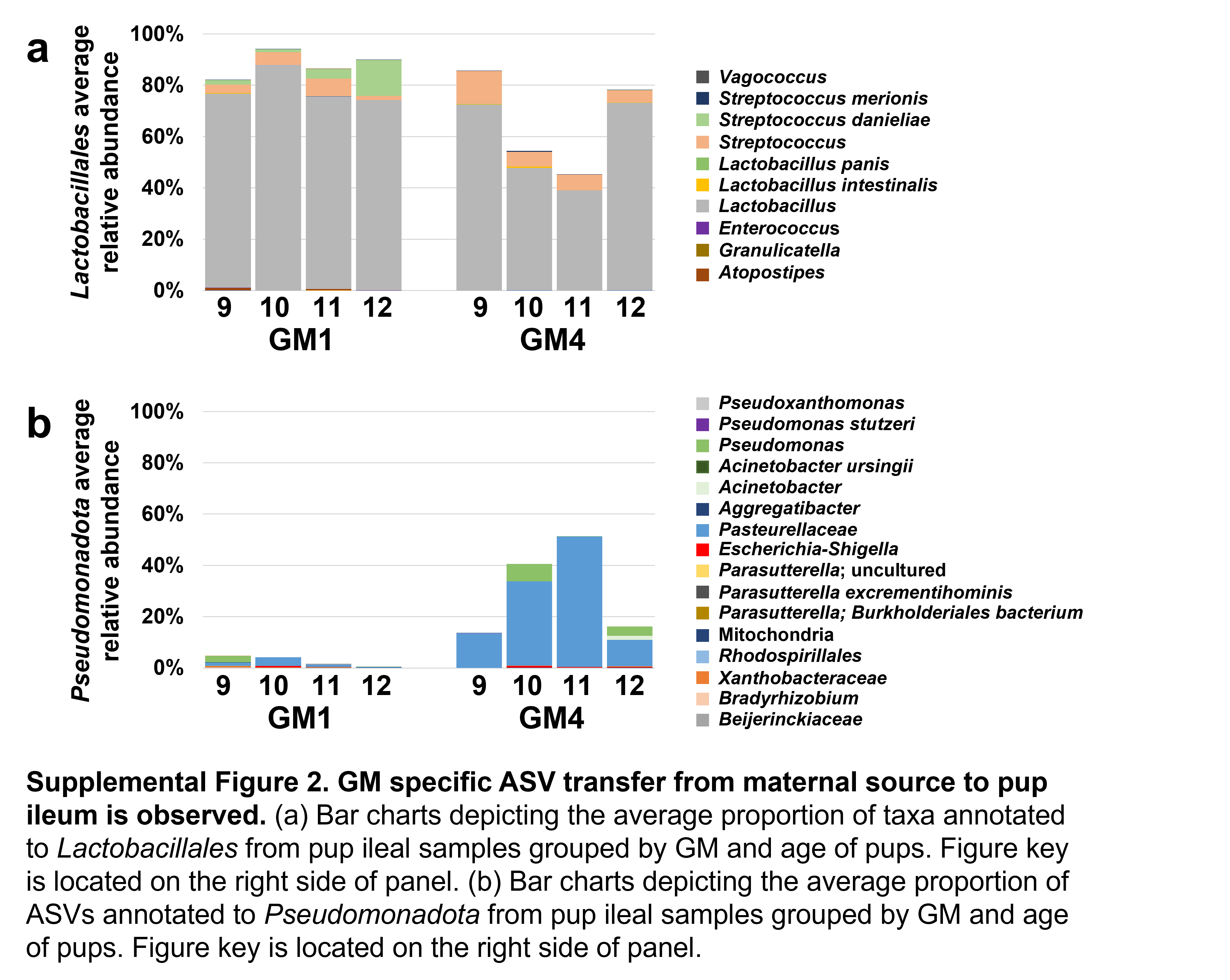

### Supplemental Figure 3

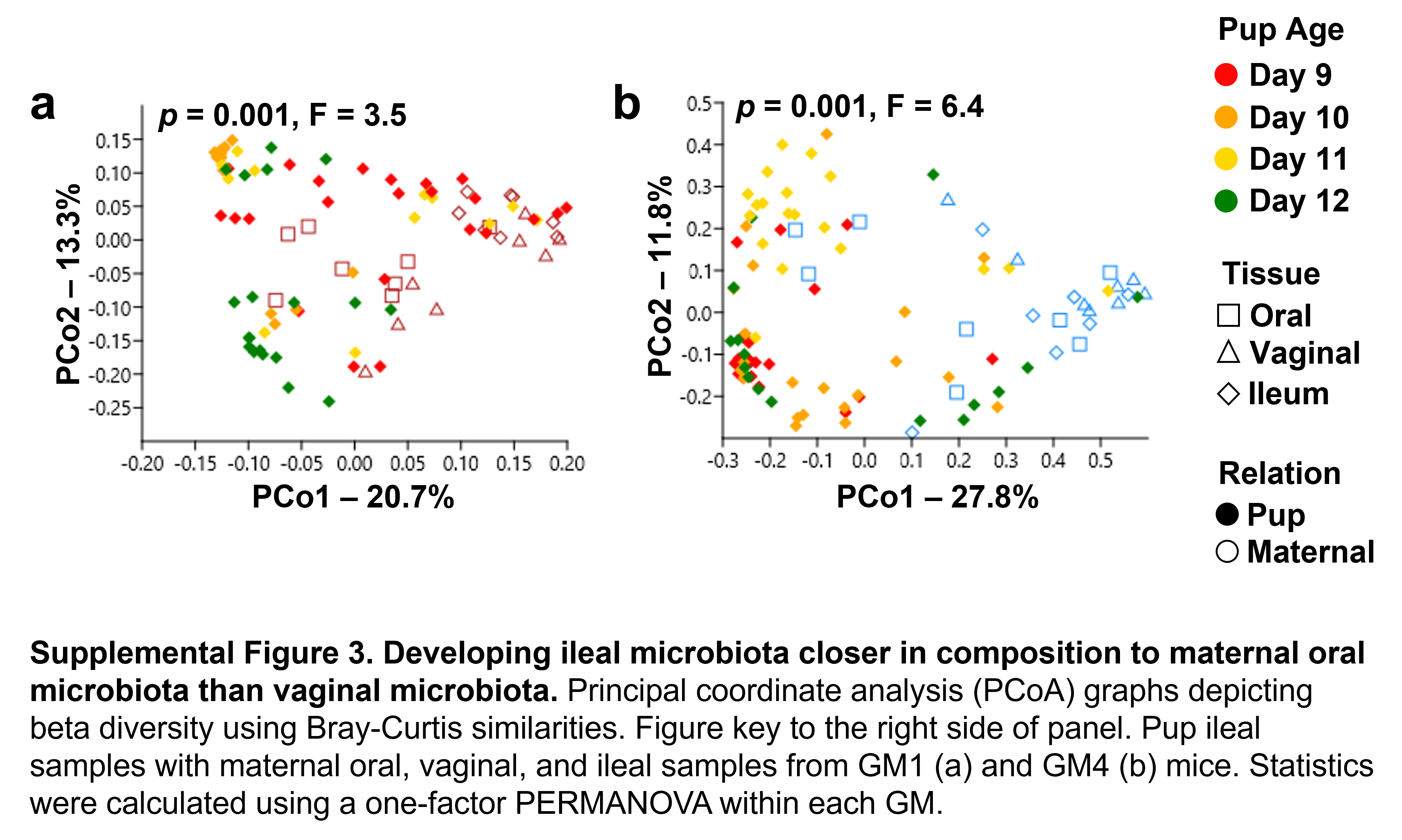

### Supplemental Figure 4

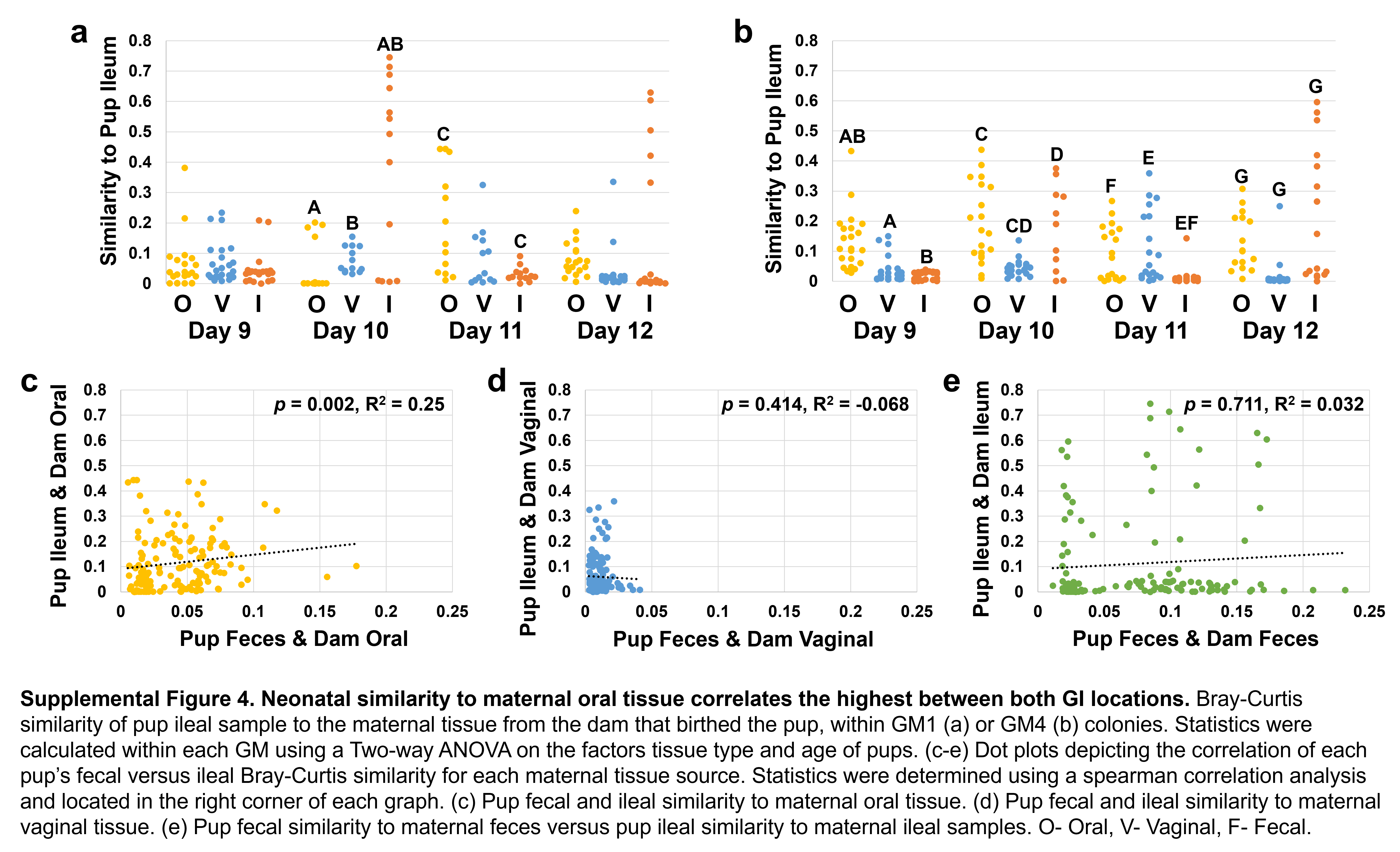

### Supplemental Figure 5

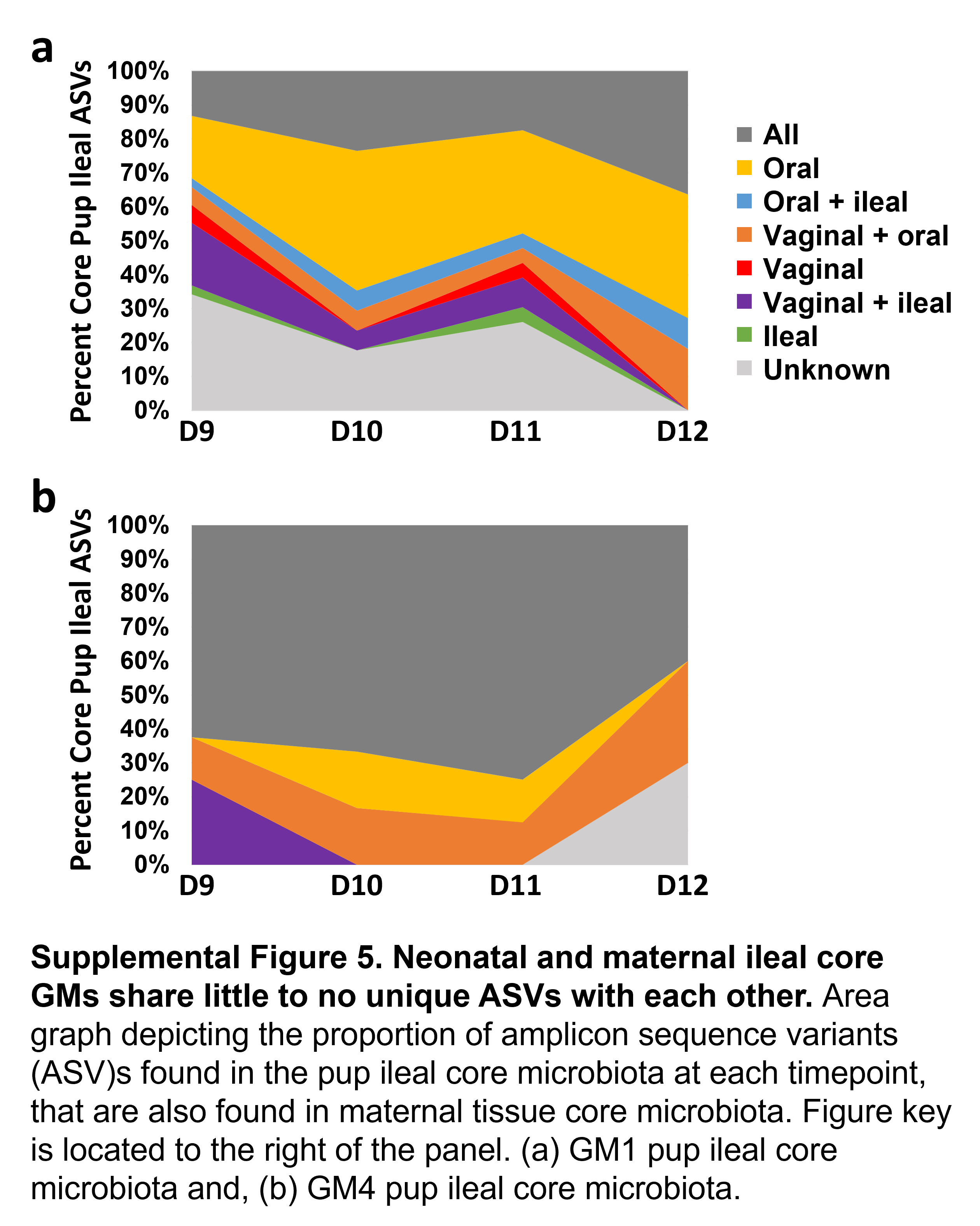
